# A novel epifluorescence microscope design and software package to record naturalistic behaviour and cell activity in freely moving *Caenorhabditis elegans*

**DOI:** 10.1101/2025.03.21.644605

**Authors:** Sebastian N. Wittekindt, Hannah Owens, Lennard Wittekindt, Aurélie Guisnet, Michael Hendricks

## Abstract

Understanding how neural circuits drive behavior requires imaging methods that capture cellular dynamics in freely moving animals. Here, we introduce Wormspy, a compact, flexible, and cost-effective microscope system paired with an open-source software package, specifically designed for high-magnification epifluorescence imaging in *Caenorhabditis elegans*. By integrating dual-channel fluorescence optics, off-the-shelf components, and a motorized stage, Wormspy enables simultaneous recording of neuronal activity and behavioral dynamics without restraining the animal. Our system incorporates both manual and automated tracking—including DeepLabCut-based feature extraction—to maintain precise centering of the subject, even during complex locomotor behaviors. We demonstrate the utility of Wormspy across multiple paradigms: quantifying body wall muscle calcium transients during locomotion, capturing rapid sensory-evoked responses in the polymodal ASH neuron during aversive stimuli, and resolving food-related activity in the AWC^ON^ neuron. Notably, Wormspy further distinguishes subcellular calcium events in the RIA axonal compartments that correlate with head bending kinematics. This versatile platform not only reproduces known phenotypes, such as altered gait in *gar-3* mutants, but also uncovers nuanced sensorimotor correlations previously inaccessible with conventional methods. Wormspy’s modular design and open-source framework lower the technical and financial barriers to high-resolution, behaviorally relevant neural imaging. Our findings establish Wormspy as a robust tool for dissecting the neural underpinnings of behavior in freely moving organisms, with potential applications extending to other small model systems.

## Introduction

The combination of smaller and cheaper imaging technology and tractable model organisms like *Caenorhabditis elegans* promises to be an important driver in our understanding of the neural correlates of behaviour. *C. elegans* is an excellent model organism for imaging-based experiments due to its small size, transparency, and genetic tools^1^. These properties have led *C. elegans* to be widely used in studies of development, cellular functioning, and neural signaling^2^, as well as being crucial for phenotyping models of disease and drug development^3^. Its cell lineage is largely invariant, allowing for individual cells to be tracked optically in both wild-type and mutant animals using differential interference contrast microscopy or fluorescent proteins^4^. Specifically, epifluorescence microscopy is a powerful tool for visualizing and studying cell activity, development, and the functioning of the nervous system.

Recent developments in hardware, such as smaller, more sensitive digital cameras, and software, such as segmentation methods that use convolutional neural networks, are increasingly making it possible to image the functioning of single cells, cell assemblies, and even whole nervous systems in freely moving animals^5^. Imaging in freely moving animals as opposed to restrained animals comes with a spate of engineering challenges, but makes possible new investigations of sensorimotor integration, navigational strategies, and a host of other behavioural research^6,7^. Existing imaging platforms developed to image freely moving *C. elegans* are broadly divided into two categories^8^. Some trackers use a high-resolution camera to image a large area and achieve higher resolution of behavioural and locomotion data by using machine vision software that can segment and track single or multiple animals^8–11^. These imaging platforms are used almost exclusively for behaviour. Other imaging platforms use high magnification objectives to achieve the resolution necessary to observe cellular processes such as neuronal calcium flux, but are more limited in the behavioural repertoire they can record ^12,13^.

Systems based on spinning disk confocal microscopes using dual objectives, one for tracking posture measurements and one for volumetric imaging of fluorescent probes, have been used for whole brain imaging in *C. elegans* crawling between an agar substrate and a coverslip^14,15^. While these systems are powerful for imaging cell bodies, several classes of *C. elegans* neurons exhibit axonal calcium events, which cannot be measured by this approach. It is unclear the extent to which animals recapitulate naturalistic behaviours under these experimental conditions. These systems are also very expensive, and the resulting datasets require considerable computational expertise to analyse and interpret. Finally, to keep the worm centred in the field of view, these imaging platforms typically use XY motors that move the stage, causing sporadic acceleration forces and vibrations, which *C. elegans* are known to be highly sensitive to^16^.

We concluded that there was a need for a compact, flexible design that would enable widefield imaging (or photoactivation) of genetically targetable cells in freely moving animals without expensive hardware or complex analysis pipelines. We therefore designed a user-friendly, inexpensive microscope and software package named Wormspy. Wormspy combines single worm tracking with a motor platform that moves the microscope relative to a fixed behavioural arena. 2-channel imaging through a single objective allows for simultaneous recordings of fluorescent signals and behaviour in a single optical path.

While we have focused on calcium imaging, Wormspy can be adapted to any number of use cases with the simple exchange of filters and objectives. The modular design of the microscope allows for a high degree of customization to meet experimental needs, such as the inclusion of optogenetic photostimulation or ratiometric recording modules and can be applied to a wide range of experimental setups. To demonstrate the potential of this design to common use cases in *C. elegans* neurobiology, we use Wormspy to characterize the gait parameters and muscle activity underlying a previously described motor coordination phenotype. We show that Wormspy can resolve sensory activity by recording activity in the ASH polymodal sensory neuron as freely moving worms encounter hyperosmotic glycerol barriers. We demonstrate how Wormspy’s ability to simultaneously combine manual and automatic tracking makes it suitable for recordings in difficult environments by imaging the activity of the food-sensing AWC^ON^ neuron at a lawn border. Finally, we demonstrate the ability of Wormspy to resolve subcellular axonal calcium transients in the RIA interneuron.

### Microscope design

For a complete description, parts list, and build guide, see Supplemental Text. Wormspy consists of a 2-channel fluorescence optical pathway mounted on a motorized stage (Fig. 1A). The modular design allows LED light sources and filter sets to be changed easily. For example, the use of a dual band dichroic mirror allows either simultaneous use of two fluorophores, a single fluorophore and brightfield image, or a photostimulation channel and a recording channel. This configuration was developed principally for calcium imaging applications in single or small numbers of neurons resolvable in the XY plane. Our configuration uses 470 and 565 nm excitation and two digital cameras, one receiving the emission spectra of GCaMP (502–538 nm), and one receiving the red transmission band from the dual band dichroic. The red transmission was used as a “bright field” image to silhouette the worm against the background for tracking and behavioural analysis, but the addition of an RFP emission filter (603–678 nm) also allowed us to use this channel for ratiometry in the ASH experiment.

**Figure 1:**
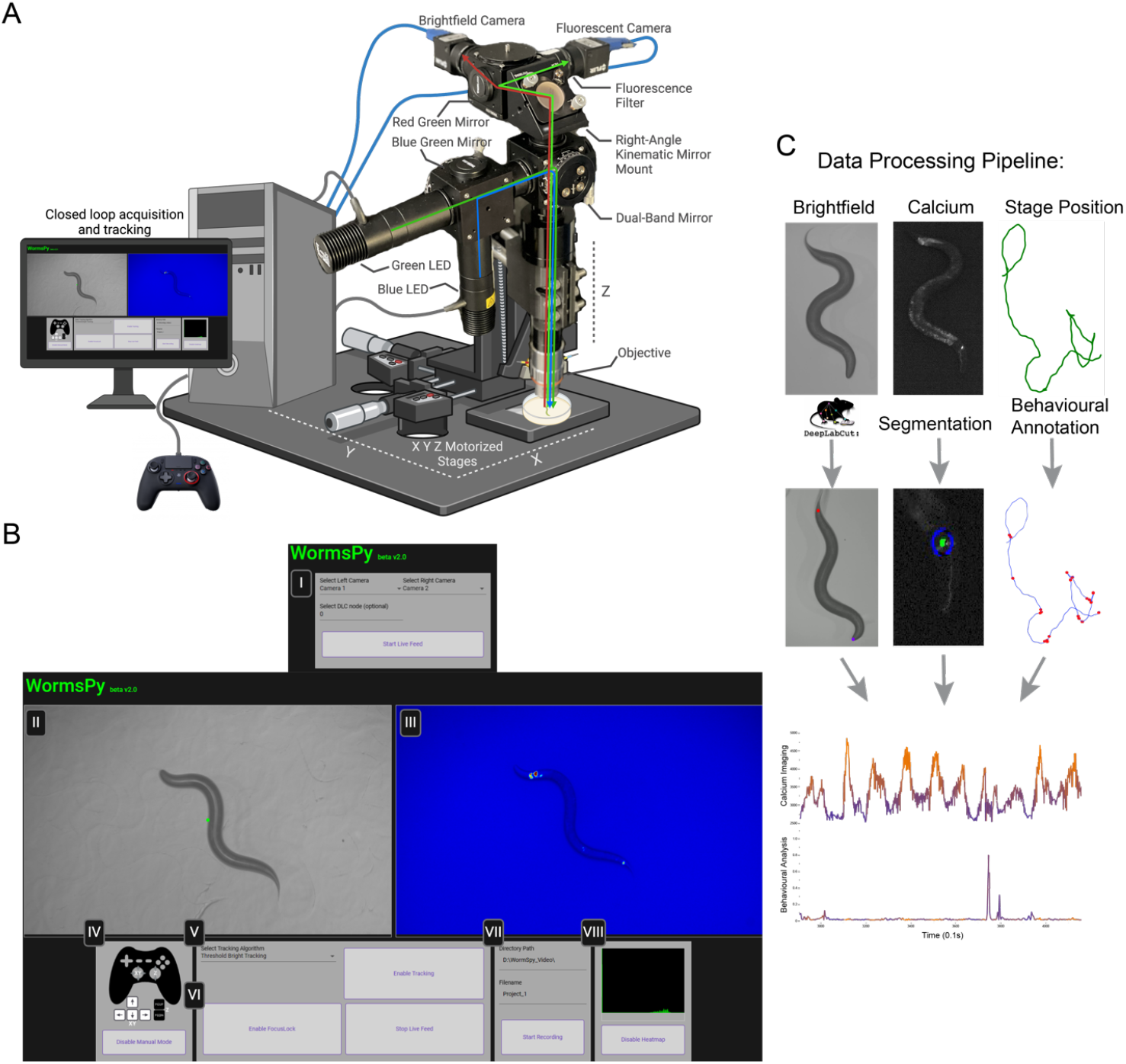
**(A)** Hardware schematic for our configuration of Wormspy. Green and Blue LEDs illuminate the stage through a dual band dichroic filter set, and the sample is captured by two cameras. XYZ motors allow for automated and manual tracking across a 50.4 x 50.4 mm area, as well as dynamic focus adjustment. **(B)** Software UI. (I) Start screen allows users to specify cameras. (II-III) Dual camera feeds. Tracking uses the left camera, whereas the right camera is recorded at 16-bit. (IV) For manual tracking a dual-joystick controller is recommended to allow for smooth adjustment of XY for tracking the animal and Z for manual focus correction. (V) Selection of tracking algorithm (VI) FocusLock adjusts the Z motor to prevent focal drift. (VII) Directory and name to save recording. (VIII) Live histogram for evaluation of fluorescent marker intensity. **(C)** Offline data processing pipeline. Brightfield videos can be further analyzed to extract pose and behavioural features. Epifluorescence intensity can be analyzed using custom segmentation code. Stage position data can be analyzed to extract behaviourally relevant information such as position in a gradient, reversals, etc. Frame-synced recordings of brightfield, calcium and stage position data enable easy comparison of behaviour and cell activity.

Light from two LEDs coupled to excitation filters (in our example, GCaMP and RFP), is directed via a dichroic mirror to a dual band filter cube and reflected onto the sample through a 100 mm tube lens and 7.5x objective lens, which provides a 1440 x 900 μm field of view, just accommodating an adult *C. elegans*. The emission path from the sample passes through the dual bandpass mirror, where it is split with another dichroic mirror and filtered into green and red components to two machine vision cameras.

While the software can be modified for any camera type, we used Teledyne FLIR computer vision USB cameras (BFS-U3-23S6M-C). We recommend selecting a sensor with a high quantum efficiency (QE) for calcium imaging, and a sensor with at least a frame rate of 10 Hz and a resolution of 1 MP for the brightfield camera (Supplemental Text).

The optical path is attached to a Z motor stage via a 90° mounting platform. We used a Zaber VSR20A vertical lift stage, which has sufficient torque to smoothly adjust the position of the microscope and keep the subject animal in focus. The Z motor in turn sits on two Zaber TSB60E horizontal translation stages that move the microscope in the XY plane to keep the subject centred in the field of view. The two cameras and the controllers for the three motors are connected via USB to a PC.

### Software design and user interface

For more details see the GitHub repository linked in the Supplemental Text. Wormspy comes with an open-source software package that includes a user interface written in Python, HTML, Typescript and open-source libraries such as OpenCV and Scikit-Image (Fig. 1B). Once both cameras are connected, the user can specify which camera corresponds to the desired left and right video feeds. If all necessary drivers are installed and the Zaber Launcher application is open to communicate with the stage motors, the program will then load the live feed of the left and right camera.

Once Wormspy is running, the user can enable *ManualMode* to use a dual-joystick controller to move the XY and Z motors to locate the animal and adjust focus. There are three automated tracking modes: 1) thresholding a dark object on a light background, which is most suitable for bright field channels; 2) thresholding fluorescent markers, which tracks the brightest object against a dark background; or 3) DeepLabCut^17^ (DLC) integration, which allows users to provide their own pre-trained model for tracking a specific feature. Each tracking method calculates the centroid—or feature of interest in the case of DLC—of the thresholded object and makes the necessary conversions for the XY motors to adjust the field of view. This conversion keeps the region of interest centred as the animal moves. On a 100 mm plate, we can track animals for >30 minutes uninterrupted. For consistent tracking we recommend a minimum frame rate of 5–10 Hz.

Once the animal has been acquired and in focus, the user can also toggle *FocusLock*, which uses the variance of the Laplacian (2^nd^ spatial derivative) to calculate the sharpness of the image from the right camera, and a proportional–integral–derivative (PID) control algorithm to smoothly adjust the Z motor to maintain the desired focus.

The user can specify a project title and directory to save the video recordings. Both cameras can save videos either as ‘MJPG’ compressed video in an audio-video interleaved (AVI) container, which is useful for behaviour, or as a lossless 16-bit tagged image file format (TIFF) stack that preserves image detail and dynamic range. Cameras are frame synced, meaning the two recordings can be directly compared by frame number. A text log of stage position and real coordinates of tracked objects is output for the duration of the recording.

## Results

### Calcium Imaging in Body Wall Muscle Cells during *C. elegans* locomotion

As our first proof of concept, we used Wormspy to record locomotion and muscle activation dynamics in wild type and uncoordinated mutants. The motor networks that produce *C. elegans* locomotion are modulated by upstream interneurons. We previously reported that disrupting the function of the RIA interneuron via mutation of *gar-3*, which encodes a muscarinic acetylcholine receptor, is associated with increased body bend amplitude^18^ (Fig. 2A).

**Figure 2:**
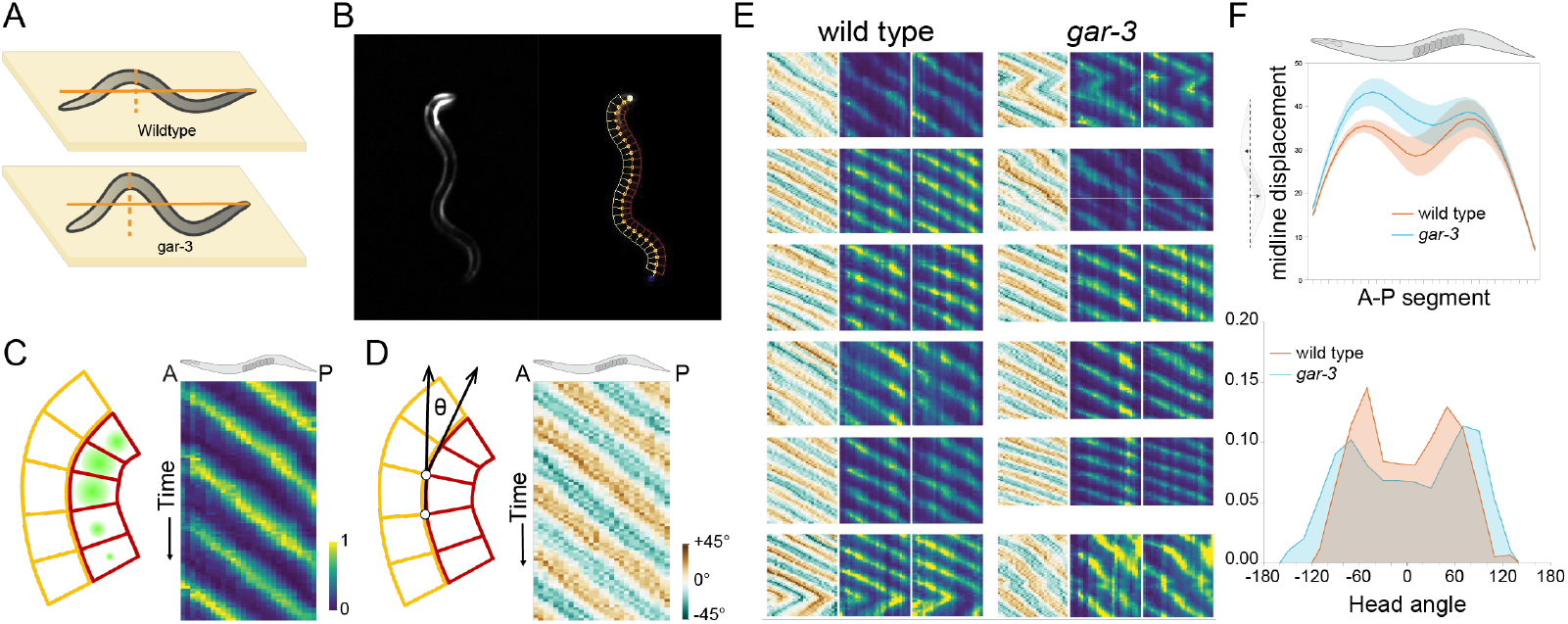
**(A)** *gar-3* displays motor coordination defects including deeper body bends. **(B)** Example frame of a calcium recording of HBR4 showing GCaMP activation in BWMCs (left) and the segmentation skeleton applied to the worm’s midline (right). **(C)** To characterize temporal and spatial distribution of calcium activity in BWMCs, worms were segmented into 28 evenly spaced ROIs by drawing perpendicular lines on either side of the worm’s midline (left). Heatmaps show calcium activity from each of these boxes arranged anterior to posterior on the X axis and temporally on the Y axis (right). **(D)** For each segment of the skeleton the angle relative to the previous segment was calculated (left) and arranged anterior to posterior as a heatmap (right). **(E)** Kymographs showing bending angle and right and left calcium activity in wildtype and *gar-3* worms. *gar-3* animals show deeper body bends and disrupted coordination. **(F)** *gar-3* worms show increased midline displacement and a wider range of head angle.

To test the ability of Wormspy to characterize the effects of *gar-3* mutation on gait and locomotion, we performed calcium imaging in freely moving animals expressing the calcium indicator GCaMP3 in body wall muscle cells (BWMCs) in wild type and *gar-3* mutants. We then analysed the videos with a custom segmentation pipeline (Fig. 2B, Supplemental Text). In each video, the animal is segmented and a midline constructed from 28 evenly space points from nose to tail, and 27 corresponding dorsal and ventral ROIs were constructed from orthogonal line segments (Fig. 2). At each segment, the midline angle relative to the anteriorly adjacent segment was calculated, and the ROIs were used to measure GCaMP intensity in opposing muscle groups in each segment. We observed the expected calcium dynamics: calcium increased in dorsal BWMCs during dorsal bends and in ventral BWMCs during ventral bends (Fig. 2C, 2D). Time series analysis for muscle contraction patterns showed that oscillations are slower in *gar-3*, corresponding to a lower gait frequency (Fig. 2E). We confirmed that *gar-3* animals displayed deeper body bends, though only in the anterior half of the body, and showed a broader distribution of head bending angles corresponding to increased gait amplitude, as previously described^18^ (Fig. 2F).

### Calcium Imaging in the Polymodal ASH Sensory Neuron

To assess Wormspy’s ability to resolve single cells and detect sensory-evoked calcium activity, we performed calcium imaging in the polymodal sensory neuron ASH as worms encountered a glycerol barrier. ASH is a ciliated neuron whose dendrite extends to the tip of the nose, where it detects noxious sensory stimuli such as harsh touch, high osmolarity, and aversive odours. Calcium activity in ASH increases rapidly following exposure to these nociceptive cues, enabling reflex-like escape behaviour including reversals^19,20^.

We adapted a behavioural assay by Ghosh et al.^21^ to measure the calcium response to a hyperosmotic glycerol barrier in freely-moving animals expressing GCaMP6f and DsRed in ASH (Fig. 3A). To control for motion artefacts, we configured Wormspy for ratiometric recording by fitting the second camera with an RFP emission filter. We tracked animals and recorded from when they approached the glycerol barrier until reversal termination. Videos of GCaMP and DsRed signals were analysed using custom segmentation code, which calculates the average of the 25 brightest pixels within the cell soma for both the red and green channel (Fig. 3C, Supplemental Text). The traces of the green and red channel were then passed through Two-channel Motion Artifact Correction^22^ to isolate GCaMP dynamics (Fig. 3B) from shared motion signals. Consistent with previous reports^21^, we found that calcium dynamics increased in the ASH soma as *C. elegans* entered the glycerol barrier in the seconds leading up to a reversal, peaking in the initial phase of the reversal (Fig. 3D).

**Figure 3:**
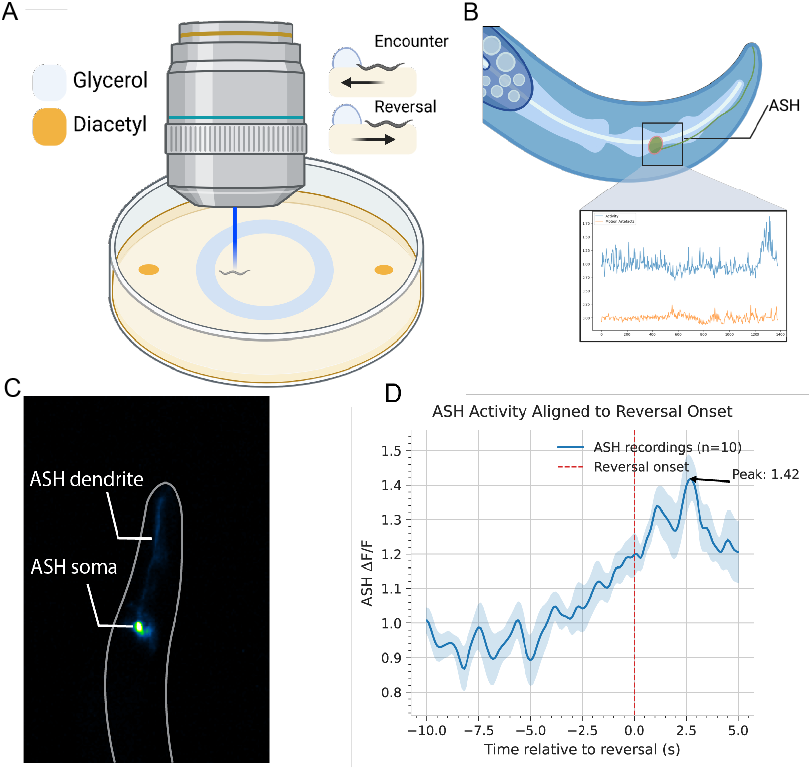
**(A)** Illustration of the ASH glycerol barrier assay. **(B)** Ratiometric recording of GCaMP6f and RFP allows for the separation of cellular activity from motion artefacts. **(C)** Example frame of TQ5856 (ASH GCaMP6f) showing the ASH cell soma and dendrite **(D)** ASH cell soma fluorescence changes aligned to reversal initiation show sustained activity increases at timescales consistent with the literature in restrained animals.

### Calcium Imaging in the AWC^ON^ Sensory Neuron on Food

Tracking and recording calcium activity presents additional challenges when *C. elegans* are on food due to variations in lighting, background, and focal depth. Wormspy’s ability to simultaneously combine manual and automatic tracking modes makes it particularly suited to this task. We tested this capability by recording calcium activity in the AWC^ON^ neuron as worms exited a patch of food.

The AWC^ON^ neuron responds to decreases in the concentration of attractive, food-associated chemicals, driving local search behavior^23^. We therefore expected that we should see an increase in AWC^ON^ calcium activity as animals exit an *E. coli* lawn. Worms were placed on a small patch of food in the middle of the plate, which was aligned with the midpoint of the Wormspy field of view. This allowed us to convert the relative position of stage movements into the worm’s absolute position in the environment (Fig. 4A, 4C). Our recordings show AWC^ON^ activity increasing concurrent with the worm’s nose leaving the patch of food, consistent with the existing literature^24^ (Fig. 4B, D).

**Figure 4:**
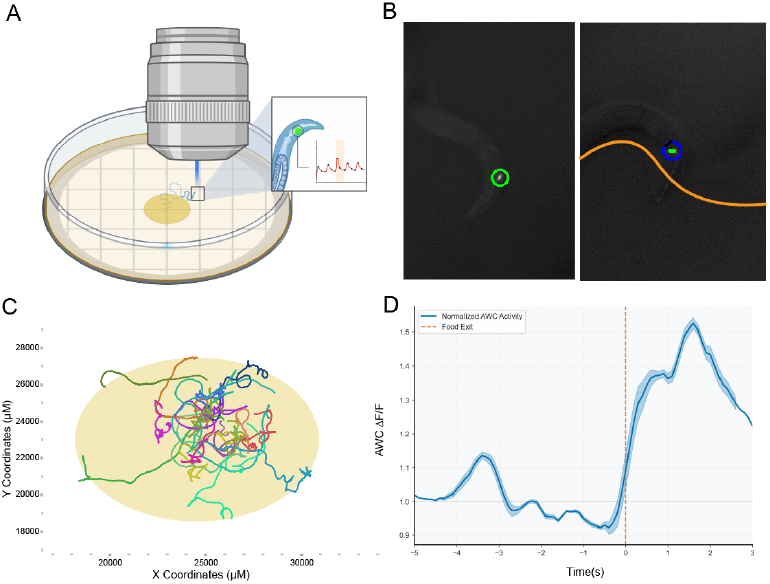
**(A)** Illustration of the AWC food patch assay. **(B)** Stage position traces showing worms navigating the food patch assay over a period of 5 minutes. **(C)** Example segmentation frames of MMH214 worms expressing GCaMP5 in AWC^ON^ navigating on food (left) and with their nose extending out of the food patch (right). **(D)** Trace of AWC^ON^ fluorescence change aligned with the worm’s nose leaving the food patch.

**Figure 5:**
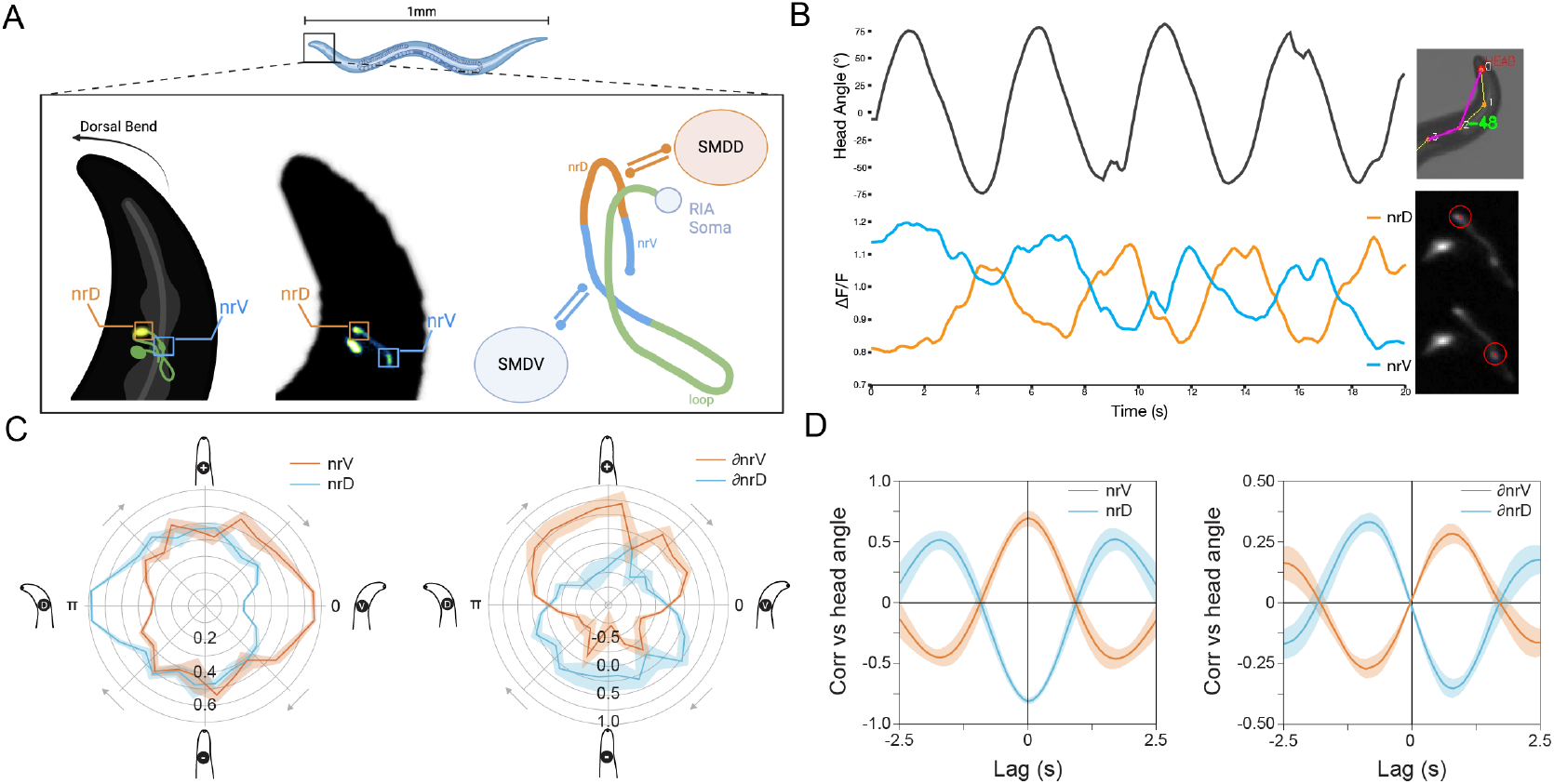
**(A)** RIA schematic: the RIA axon exhibits local calcium events in the nerve ring compartments nrV and nrD that encode head movements via reciprocal connections to SMDV and SMDD. **(B)** Example trace of head angle alongside nrV and nrD activity over time. Head angle was calculated from points placed along the midline of the worm, and nrD and nrV were segmented using a custom SAM2 analysis pipeline. **(C)** Polar plots showing nrV and nrD activity plotted according to oscillation period (left) and the time derivative of nrV and nrD (right) **(D)** Cross-Correlation of nrV and nrD signal vs head angle (left) and their time derivatives (right).

### Calcium Imaging in RIA axonal compartments

To demonstrate that Wormspy can resolve behaviourally relevant signals in subcellular compartments, we imaged worms expressing GCaMP6f in RIA. As previously described, local, compartmentalized calcium events in the ventral and dorsal segments of the RIA axon (nrV and nrD) are correlated with ventral and dorsal head bends, respectively (Fig. 4A)^18^. The orientation of the axon is such that the nrV and nrD compartments are resolvable in the same focal plane, allowing us to capture the activity in both segments of the axon as worms performed head bends during forward locomotion. Using the brightfield image, we extracted head angles for each frame by plotting points along the animal’s midline. We then used this angle to cancel out rotational movement of the neuron in the GCaMP recordings and used Segment Anything Model 2 (SAM2)^25^ to define consistent masks for both the nrV and nrD axonal compartments (Fig. 4B). We estimated the local period of head bends for each animal over time. By plotting nrV and nrD activity according to head oscillation period, we show that nrV activity peaks at the most ventral head position, and nrD peaks with dorsal head positions. Because the motor neuron activity driving local RIA calcium dynamics should be active during bending rather than at the terminus of a bend, we also plotted the derivatives of local calcium domain signals. Period analysis and cross-correlations showed that the peak rate of calcium increase in each compartment corresponds to peak ventral (positive) or dorsal (negative) head angle velocity. Interestingly, our previous measurements of RIA calcium in restrained worms showed a slightly different periodicity in which peak calcium levels and rates of increase lagged peak head position and velocity. This may reflect the fact that when restrained in microfluidic devices, animals rarely exhibit the regular head oscillations of forward locomotion and spend a large percentage of time in a reversal state, which may involve different timing of motor commands.

## Discussion

Recording and analysing behavioural and neural activity is at the core of neuroscience research. The potential exists for this technology to be made much more widely available through low-cost, open-source systems. We believe our microscope design is a contribution to this effort. Robust recording of epifluorescence signals and cellular activity has long been difficult in imaging environments that do not incapacitate or restrain the animal. Our design is intended to make fluorescence imaging in freely moving animals as inexpensive and simple as possible, without relying on expensive commercial microscopy systems. While wide field imaging does not allow optical sectioning or resolving cells in the Z-axis, it is more robust with respect to focal movement artifacts, thus common approaches that impede natural movement, like compressing animals between a coverslip and the agar surface are not necessary.

Wormspy was able to resolve the phasic calcium transients of BWMCs and track worms for long intervals to characterize locomotion phenotypes. We imaged the AWC^ON^ neuron as worms exited a patch of food and demonstrated the platform’s ability to resolve single neurons in complex imaging environments. Given that both cameras use the same light path and frame-sync recordings, users can align the images in post. Analysis of both the behavioural data and neural activity collected using Wormspy allows users to make inferences about how neural activity affects behaviour and how the worm’s interactions with its environment affect neural activity.

Additionally, for more motion sensitive applications, we demonstrated Wormspy’s potential for ratiometric recording by imaging the ASH sensory neurons as worms reversed to escape a glycerol barrier. In RIA we show that it is possible to record calcium activity not only in neuronal cell bodies, but also in dendritic or axonal compartments, provided a sufficiently bright GCaMP line is used. For example, in the RIA::GCaMP6f strain used here, we were able to record with an exposure time of 14 ms, minimizing motion artefacts. Our system allowed us to show for the first time how localized calcium dynamics in the RIA axon are correlated with locomotion, identifying temporal differences from data collected in restrained animals.

While Wormspy was developed for calcium imaging in head sensory and interneurons in *C. elegans*, its modular design and low cost allows it to be iterated on by other researchers and find applications in other fields. Our system is built with off-the-shelf components and is intended to be highly extensible and modifiable. It is particularly amenable to applications like closed-loop optogenetic stimulation. We anticipate that it can easily be applied to other small organisms such as *Drosophila melanogaster* larvae (see recordings in Supplemental Text).

## Materials and methods

### Animals

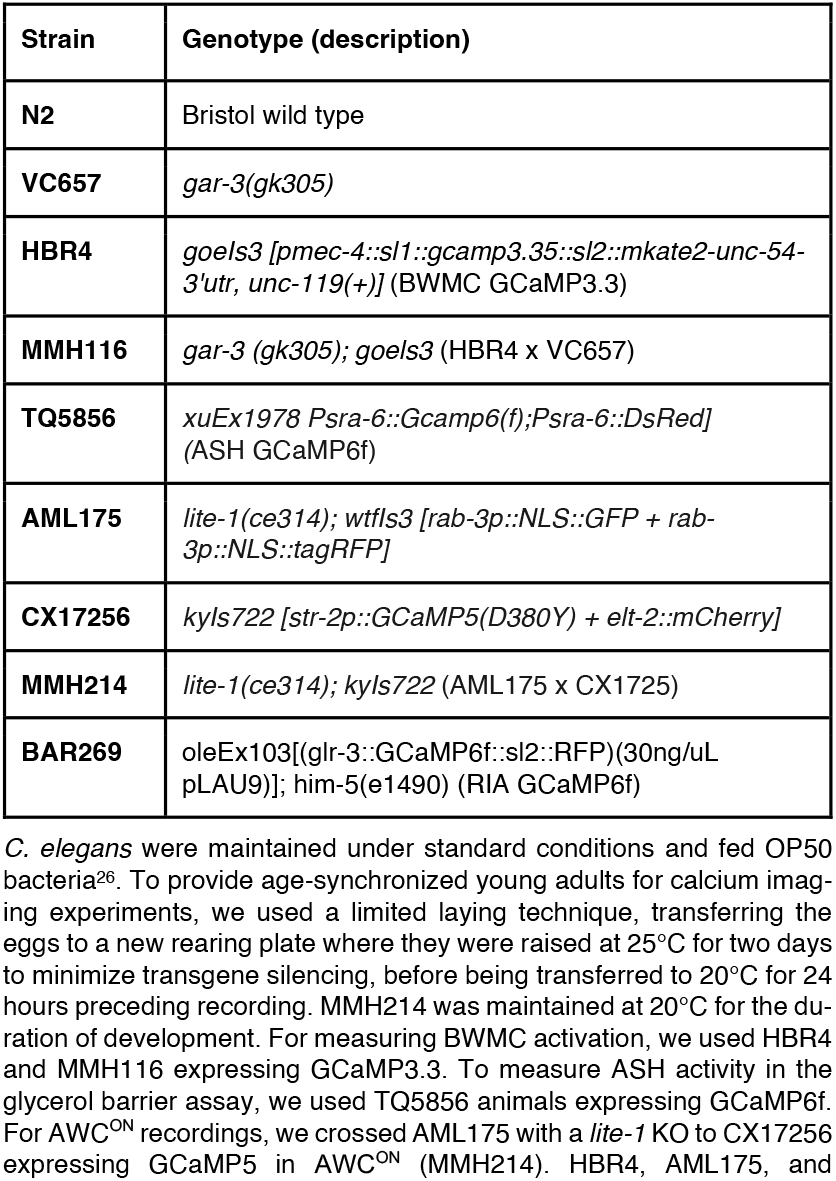
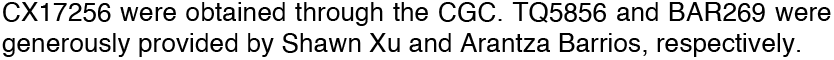

*C. elegans* were maintained under standard conditions and fed OP50 bacteria^26^. To provide age-synchronized young adults for calcium imaging experiments, we used a limited laying technique, transferring the eggs to a new rearing plate where they were raised at 25°C for two days to minimize transgene silencing, before being transferred to 20°C for 24 hours preceding recording. MMH214 was maintained at 20°C for the duration of development. For measuring BWMC activation, we used HBR4 and MMH116 expressing GCaMP3.3. To measure ASH activity in the glycerol barrier assay, we used TQ5856 animals expressing GCaMP6f. For AWC^ON^ recordings, we crossed AML175 with a *lite-1* KO to CX17256 expressing GCaMP5 in AWC^ON^ (MMH214). HBR4, AML175, and CX17256 were obtained through the CGC. TQ5856 and BAR269 were generously provided by Shawn Xu and Arantza Barrios, respectively.

### Behaviour

#### Calcium imaging in body wall muscles

We transferred well-fed adult day 1 worms (HBR4, MMH116) individually to medium unseeded NGM plates before placing them under the Wormspy objective. The worms were illuminated with blue light and tracked using the user interface. Worms were recorded as they moved across the agar for five minutes. We then analysed the videos with a custom segmentation pipeline to measure ventral and dorsal BWMC activity (Fig. 2B, Supplemental Text).

#### Calcium imaging in the ASH neurons

Young adult TQ5856 worms were starved for 30 minutes. A ring of glycerol (≥99.0%, Sigma-Aldrich) two centimetres in diameter (approximately 12 μL in volume) was pipetted in the centre of medium unseeded NGM plates. To incentivize animals to enter the aversive glycerol barrier, two 1 uL drops of diacetyl (2,3-Butanedione, Sigma-Aldrich, 1:350 dilution) were pipetted outside of the ring, at opposite ends of the plate, about 1 cm from the border^18^ (Fig. 3A). The glycerol was allowed to absorb into the agar for ten minutes. Individual animals were then placed inside the centre of the ring and left to crawl freely for five minutes while being tracked and recorded using Wormspy. Blue light was turned on as worms approached the barrier to minimize bleaching. Changes of fluorescence intensity of the calcium indicator GCaMP6f in the ASH neuron were recorded via the GFP-filtered camera. Recordings where the worms encounter the glycerol barrier were segmented for the ASH cell soma via a custom segmentation pipeline (Fig. 3B,C, Supplemental Text). Frame counts where animals entered the glycerol barrier and initiated reversal were manually ascertained from the brightfield recordings.

#### Calcium imaging in the AWC^ON^ neuron

Well-fed MMH214 worms were individually transferred onto small NGM plates with a small patch of food in the centre. Using a grid visible underneath the NGM plates, the plates were aligned such that at the start of a recording the worm in the patch of food was as close as possible to the midpoint of the Wormspy range of motion (home position). Using a combination of automatic and manual tracking, worms were kept in view and in focus for 5 minutes as they explored the patch of food. Recordings were manually analysed to identify frames where the worm’s nose exited the food. The brightest 25 pixels of the AWC^ON^ cell soma in the GCaMP6f channel was measured for each frame.

#### Calcium imaging in RIA axonal compartments

Well-fed BAR269 worms were individually transferred onto medium agarose plates without food and recorded using automatic tracking for 1-2 minutes at a framerate of 10 Hz and 14 ms exposure time to minimize motion artefacts. For each worm, 200 frames of uninterrupted forward runs were selected for analysis. The worm in the brightfield image was skeletonized by placing 12 evenly spaced points along the midline. Head angle was calculated using the angle between the vectors corresponding to points 3→2 and 2→0. The GCaMP6f channel recordings were cropped into 80×80px frames centred on the neuron soma. Using custom code, we first used head orientation from the brightfield channel to rotationally register each neuron and then prompted the Segment Anything Model 2 (SAM2) to recognize the nrV and nrD compartments and create custom masks that were robust throughout the recording. The brightest 25 pixels in each mask were used for analysis. The traces were smoothed with an equally weighted moving average of window size 0.5 seconds (5 frames). nrV and nrD calcium were min/max normalized (0–1), and head angles were normalized to the deepest dorsal (−1) or ventral (+1) bend. Derivatives were calculated between adjacent time points on smoothed, normalized values. To estimate the period of oscillation a phase angle was calculated as *ϕ=atan2(∂angle,angle)*. Polar plots show mean values within 24 15° bins. For figures 4C and 4D binned by polar coordinates and averaged across worms (n=7).

## Author contributions

S.W. developed the Wormspy hardware platform. L.W. contributed to the software development together with S.W. S.W. and H.O. designed experiments and collected data. S.W., H.O., A.G., and M.H. analyzed data and wrote the manuscript. M.H. obtained funding.

## Acknowledgments

This study was funded by the Canadian Institutes of Health Research (CIHR PJT-155980), Canadian Foundation for Innovation (CFI) (32581), the Canada Research Chairs Program (950-231541) and McGill University. H.O. is supported by a National Sciences and Engineering Research Council PGS-D doctoral fellowship (PGSD-579692). We thank Arantza Barrios and Shawn Xu for strains. Elliott Hendricks assisted in the early stages of software development. A special thank you to Dr. Arjun Krishnaswamy for advice on instrument design. Some strains were provided by the CGC, which is funded by NIH Office of Research Infrastructure Programs (P40 OD010440). Typeset with the bioRxiv word template by Chrelli: www.github.com/chrelli/bioRxiv-word-template.

## Supplemental Text: Build guide and links

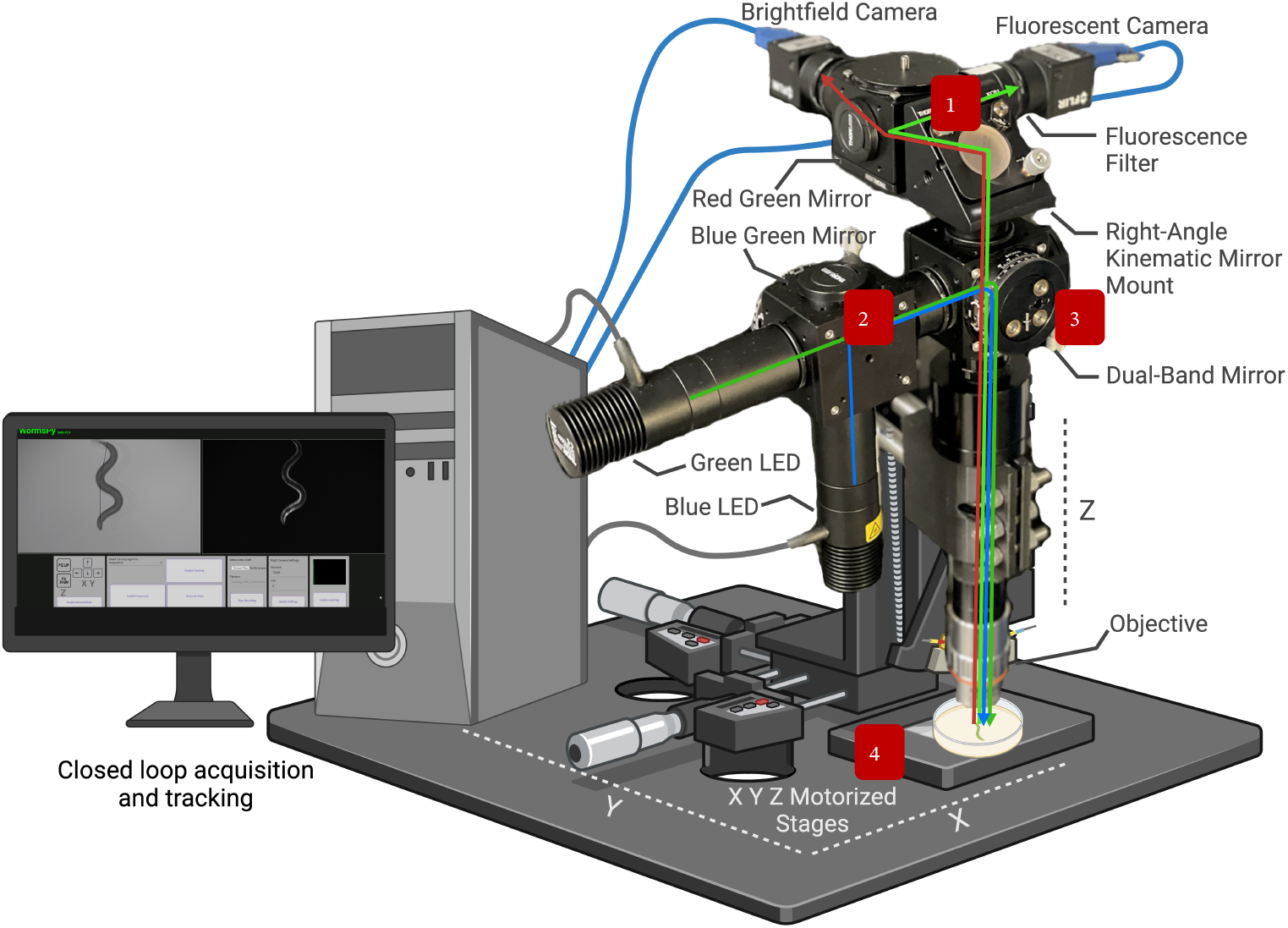

### Wormspy Dual Camera Build Guide

Overview: *[#] refers to the index number of the component in the table below.

The core of the build is three ThorLabs 30mm cage cubes [1] that have a blank cover plate [2] on one side and the mirrors on the other side, mounted into the cage cubes via precision rotation or fixed platforms [3,4,5]. Components are connected to the cage cubes via adjustable ThorLabs threaded tubes as needed [6]. Our build is designed for ratiometric calcium imaging at high magnifications and this guide will use the components we selected for that purpose, but they can easily be exchanged or modified as needed.

1. Imaging Cube:
  1. The imaging cube consists of two computer vision cameras [7] selected to have high quantum efficiency for maximum sensitivity to phasic calcium transients. Red and green emission filters [8,9] are mounted on the inside of a short lens tube [10] just before the camera. The bandwidth of emitted light is split via a red/green mirror [11] mounted on a fixed cage cube platform [4,5] as we found this to be sufficient to align the cameras. This can also be swapped out for a precision rotation platform [3,5]. The imaging cube is connected to the light path cube via a right-angle kinematic mirror mount for X Y alignment [12,13].
2. LED Excitation Cube:
  1. The LED excitation cube consists of two single-colour mounted LEDs at 470nm and 565nm [14,15] driven by an LED Driver [16] and filtered by excitation filters selected for GCaMP/mCherry ratiometry [17,18]. The LEDs are diffused via an aspheric condenser lens [19] mounted in an SM30 Lens Tube [20] with a retaining ring [21]. A blue/green mirror [22] mounted on a precision rotation platform [3,5] directs both LEDs onto the central light-path cube.
3. Light-Path Cube:
  1. The light-path cube is connected to the other two cubes via adjustable c-mount extension tubes [6] and houses a dual-band dichroic mirror [23] mounted on a precision rotation platform [3,5]. The bottom of the cube is attached to a tube lens [24], a 100mm lens tube [25] and an infinity-corrected objective [26] that focuses the light on the sample.
4. Motor Platforms:
  1. The entire imaging platform is attached to the motors via a 3D printed clamp [29] (commercial clamps are also available) that attaches to the 100mm lens tube [25]. The clamp is in turn screwed onto two optical posts [28] that attach to the right-angle mounting plate [27] that is attached to the top of the vertical lift stage [30,31,32,33] that controls the Z axis. The vertical lift stage is mounted on top of two TSB translational stages [34,35,36] that control the X and Y axis. The motors and cameras are connected via USB to a PC. If the correct Zaber motion and Spinnaker drivers are installed, this setup should be compatible with Wormspy.

Note: C-mount connectors of various lengths were used as needed to ensure components were aligned with one another. The use of a tube lens ensures the entire light path remains in focus, regardless of length. However, an overall light path > 200mm will change the size of the focused image on the camera sensor. The ThorLabs YouTube channel is a good resource for determining optimal light path lengths for your components.

PARTS LIST:

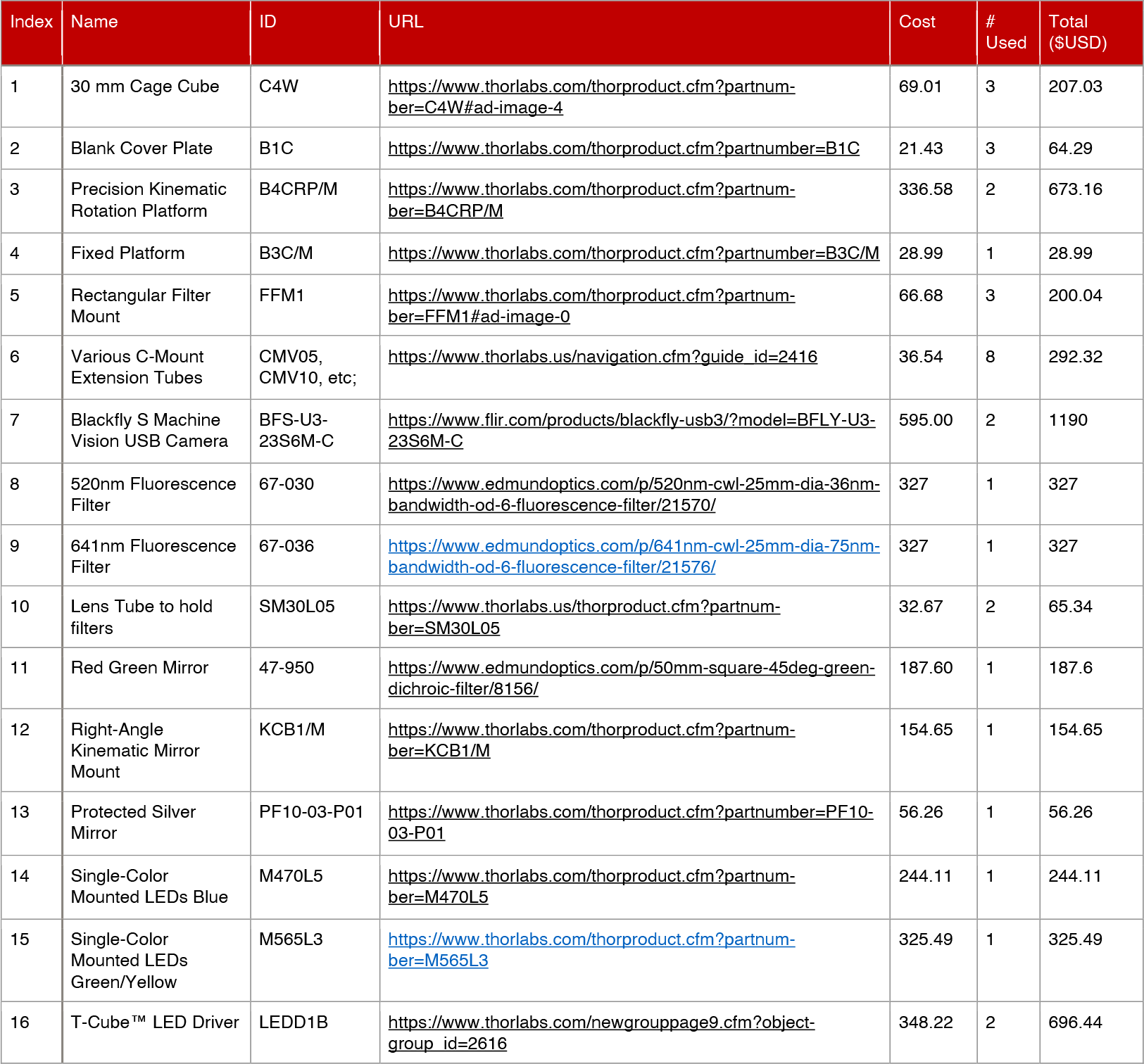
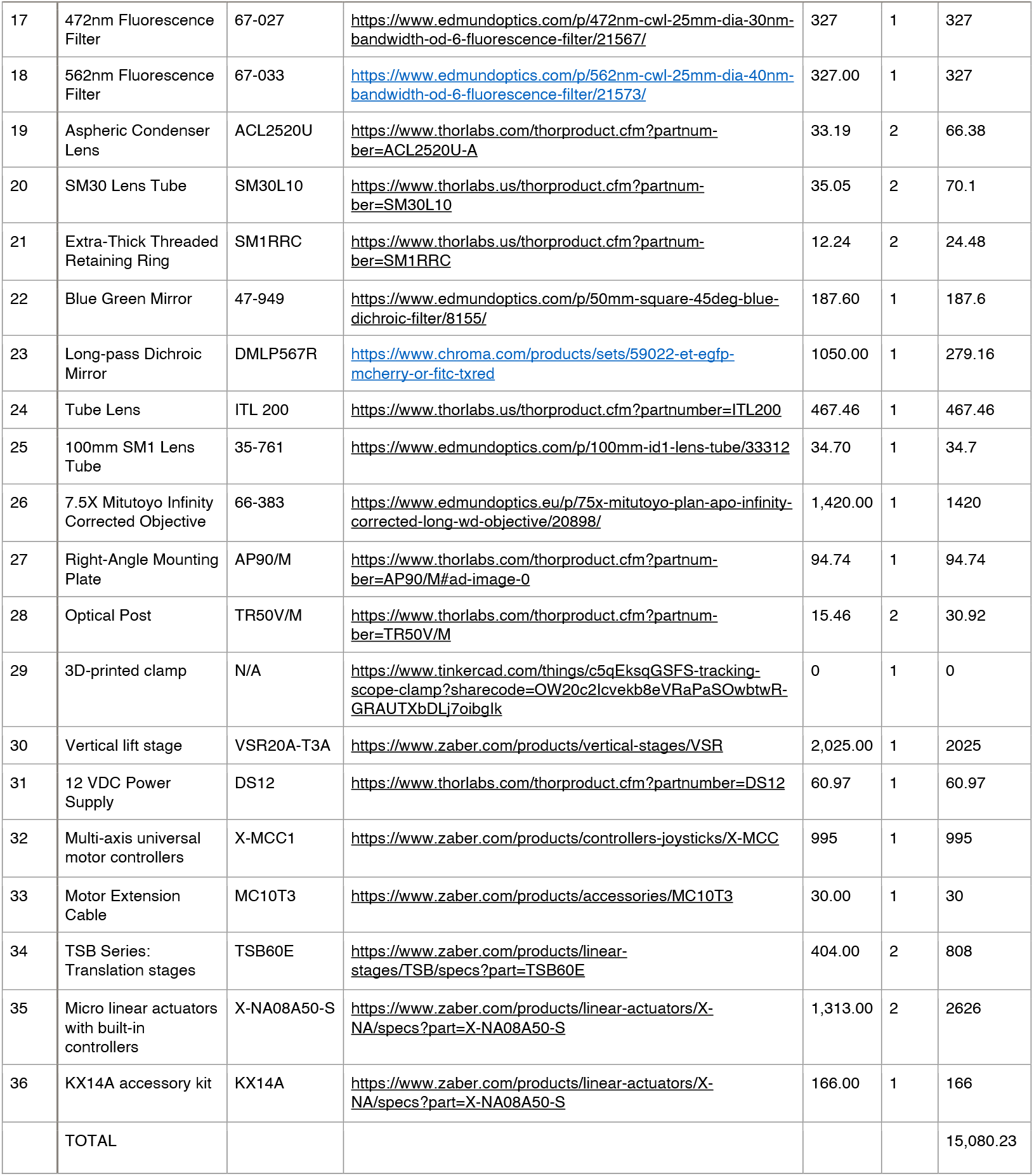

## Links to videos and code

### Wormspy GitHub

Wormspy (sebzdead.github.io) -- Includes installation and setup instructions

### User Interface Demonstration Video

https://youtu.be/mhyYDpziSE8 -- demonstrates the workflow

### Segmentation and analysis code

https://github.com/Hendricks-Worm-Lab/WormsPy_paper_analysis

### Using Wormspy to record GCaMP expressed in Drosophila larvae muscle

https://sebzdead.github.io/WormsPy/media/drosophila.gif

